# Elucidation of the 14-3-3ζ interactome reveals critical roles of RNA splicing factors during adipogenesis

**DOI:** 10.1101/184499

**Authors:** Yves Mugabo, Mina Sadeghi, Nancy N. Fang, Thibault Mayor, Gareth E. Lim

**Affiliations:** CRCHUM, Montréal, QC H2X 029, Canada; Department of Medicine, Université de Montréal, Montréal, QC H3T 1J4, Canada; Department of Biochemistry and Molecular Biology, University of British Columbia. Vancouver, Canada; Present Address: Department of Medical Genetics, Harvard Medical School, Boston, USA.

**Keywords:** 14-3-3, adipocyte, RNA splicing, proteomics, adipogenesis, interactome

## Abstract

Adipogenesis is facilitated by a complex signaling network requiring strict temporal and spatial organization of effector molecules. Molecular scaffolds, such as 14-3-3 proteins, coordinate such events, and we have previously identified 14-3-3ζ as an essential scaffold in adipocyte differentiation. The interactome of 14-3-3ζ is large and diverse, and it is possible that novel adipogenic factors may be present within it. Mouse embryonic fibroblasts from mice over-expressing a TAP-epitope-tagged 14-3-3ζ molecule were generated, and following the induction of adipogenesis, TAP-14-3-3ζ complexes were purified, followed by mass spectrometry analysis to determine the 14-3-3ζ interactome. Over 100 proteins were identified as being unique to adipocyte differentiation, of which 56 were novel interacting partners. Previously established regulators of adipogenesis (ie, Ptrf/Cavin1 and Phb2) were found within the 14-3-3ζ interactome, confirming the ability of this approach to identify regulators of adipocyte differentiation. An enrichment of proteins in the interactome related to RNA metabolism, processing, and splicing was identified, and analysis of transcriptomic data revealed that 14-3-3ζ depletion in 3T3-L1 cells affected the alternative splicing of mRNA during adipocyte differentiation. Of the RNA splicing factors within the 14-3-3ζ interactome, depletion of Hnrnpf, Hnrnpk, Ddx6, and Sfpq by siRNA revealed essential roles of these proteins in adipogenesis and their roles in the alternative splicing of *Lpin1*. In summary, novel adipogenic factors can be detected within the 14-3-3ζ interactome, and further characterization of additional proteins within the 14-3-3ζ interactome has the potential of identifying novel targets to block the expansion of adipose tissue mass that occurs in obesity.

## 1. Introduction

Central to the development of obesity are the increases in number and size of adipocytes according to nutrient availability (1,2). Despite various therapies to limit weight gain and promote weight loss, it is surprising that none specifically target the adipocyte to limit its expansion or growth (1,2). The complex transcriptional network and cellular processes that govern the differentiation of adipocyte progenitor cells contributes to the difficulty in targeting adipocytes therapeutically (1,2). Protein phosphorylation is a key post-translational modification that determines the activation state, subcellular localization, and stability of adipogenic regulators (3–7). Furthermore, phosphorylation status also determines their interactions with molecular scaffold proteins, which aid in the coordination of complex transcriptional networks (3,4).

We previously identified the molecular scaffold, 14-3-3ζ, as a critical regulator of glucose homeostasis and adipogenesis (4,8,9). Specific to the adipocyte, systemic deletion of 14-3-3ζ in mice significantly reduced visceral adiposity and impaired adipocyte differentiation, whereas transgenic over-expression of 14-3-3ζ exacerbated high-fat diet induced obesity (4). The hedgehog transcription factor, Gli3, was identified as a critical downstream effector in 14-3-3ζ-mediated adipogenesis, but the diversity of proteins in the 14-3-3ζ interactome suggest the possibility that other interacting proteins or pathways parallel to Gli3 may be also involved.

Unbiased approaches, such as proteomics and transcriptomics, can lead to the discovery of novel factors that drive adipogenesis, in addition to providing insight into physiological pathways influenced by adipogenic regulators like 14-3-3ζ (4,10–15). All seven mammalian 14-3-3 isoforms have large, diverse interactomes (8,16–18), and they are dynamic and change in response to various stimuli (11–13,19). Thus, inducing pre-adipocytes to differentiate may permit the identification of novel differentiation-specific factors within the 14-3-3ζ interactome and reveal pathways and biological processes that are essential to the development of a mature adipocyte.

To elucidate the 14-3-3ζ interactome during adipogenesis, we employed a proteomic-based discovery approach. Herein, we report that previously established factors required for adipogenesis (ie, Ptrf/Cavin1 and Phb2) can be detected in the interactome, and novel factors, such as those involved in RNA splicing, are also enriched in the interactome during differentiation. To test for their roles in adipogenesis, siRNA knockdown approaches were used and revealed the requirement for RNA splicing factors, such as Hnrnpf, Sfpq, and Ddx6. Taken together these findings demonstrate the usefulness of examining the interactome of 14-3-3 proteins in the context of a physiological process, such as adipocyte differentiation, and highlight the ability to find novel functional regulators through this approach. Understanding how the interactome is influenced by disease states, such as obesity, may lead to the identification of novel proteins that contribute to disease pathogenesis.

## 2. Material and methods

### 2.1 Generation of 14-3-3ζ^TAP^ MEFs and Cell culture

Embryos at e13.5 were harvested from pregnant transgenic mice over-expressing a TAP-epitope-tagged 14-3-3ζ molecule (4), and mouse embryonic fibroblasts (MEFs) were generated according to established protocols. 3T3-L1 cells (between passages 11-17) and mouse embryonic fibroblasts (MEFs) were maintained in 25mM glucose DMEM, supplemented with 10% newborn calf serum or fetal bovine serum (FBS), respectively, and 1% penicillin/streptomycin (ThermoFisher Scientific, Waltham, MA). Differentiation of MEFs and 3T3-L1 cells was induced with DMEM, supplemented with 10% FBS, 172 nM insulin, 500 μM IBMX, and 500 nM dexamethasone (MDI). Differentiation media for MEFs was further supplemented with rosiglitazone (Sigma-Aldrich, Oakville, ON, Canada). Following incubation with differentiation media for 2 days, media was replaced every two days with 25mM glucose DMEM, supplemented with 10% fetal bovine serum and 172 nM insulin. Differentiation was assessed by Oil Red-O incorporation (Sigma-Aldrich), as previously described (4).

### 2.2 Mass spectrometry

Equal amounts of cell lysates from undifferentiated and differentiated TAP-14-3-3ζ MEFs were subjected to an overnight incubation with IgG coupled to protein-G beads (ThermoFisher Scientific) in RIPA buffer. Bound proteins from each pull down were eluted with 1X SDS sample buffer without reducing agents and separated by SDS-PAGE prior to in-gel digestion (20). For each sample, peptides from three fractions (<50KDa, >50KDa, IgG bands) were then purified on C-18 stage tips (21) and analyzed using a LTQ-Orbitrap Velos (ThermoFisher Scientific) as previously described (22). Data were processed with Proteome Discoverer v. 1.2 (ThermoFisher Scientific) followed by a Mascot analysis (2.3.0, Matrix Science, Boston, MA) using the *Uniprot-Swissprot_mouse* protein database (05302013, 540261 protein sequences). Only proteins with at least two peptides (false positive discovery rate <=1%) in one of the two samples were retained. Two independent pull-downs were used for mass spectrometry and proteomic analysis. Proteins were analyzed with DAVID and String-Db to analyze proteins based their biological processes (23,24).

### 2.3 Analysis of differential exon usage

To understand how adipocyte differentiation and depletion of 14-3-3ζ affected alternative splicing of mRNA, differential exon usage via DEXSeq was used as a surrogate measurement (25). Our previous transcriptomic data [GSE60745] (26) were aligned to the mouse genome (Ensembl NCBIM37) via Tophat (v. 2.1.1), and the number of reads mapping to a particular exon were compared to the total number of exons in a given gene (25). A false discovery rate (FDR) of 0.05 was used to filter results. This dataset was also analyzed to examined how depletion of 14-3-3ζ or differentiation affects the expression profile of target genes. Genes identified by DEXSeq were subjected to gene ontology analysis to categorize genes by biological function (27). Alternatively, analysis of *Lpin1* splicing was performed by RT-PCR, as described previously (28). PCR products were resolved on an agarose gel, followed by densitometric analysis of splice variants by ImageJ (29).

### 2.4 siRNA-mediated knockdown, RNA isolation and quantitative PCR

3T3-L1 cells were seeded at a density of 75,000 per well prior to transfection with control siRNA or target-specific Silencer Select siRNAs (ThermoFisher Scientific). Transfection was performed using Lipofectamine RNAimax, as per manufacturer instructions (ThermoFisher Scientific), at a final siRNA concentration of 20 μM per well. RNA was isolated from 3T3-L1 adipocytes or MEFs with the RNEasy kit (Qiagen, Mississauga, ON, Canada). Synthesis of cDNA was performed with the qScript cDNA Synthesis kit (Quanta Biosciences, Gaithersburg, MD), and transcript levels were measured with SYBR green chemistry or Taqman assays on a QuantStudio 6-flex Real-time PCR System (ThermoFisher Scientific). Primer sequences are available on request. All data were normalized to HPRT by the 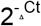 method, as previously described (4,9,30). Confirmation that knockdown of 14-3-3ζ has no effect of global RNA transcription was determined using the Click-iT RNA Alexa 488 imaging kit, as per manufacturer instructions (Thermo Scientific).

### 2.5 Statistical Analysis

All data were analyzed by one- or two-way ANOVA, followed by appropriate post-hoc tests, or by Student’s t-test. Data were considered significant when p<0.05.

## 3. Results

### 3.1 Generation of TAP-14-3-3ζ mouse embryonic fibroblasts (MEFs)

To examine how adipocyte differentiation influences the 14-3-3ζ interactome, we generated mouse embryonic fibroblasts (MEFs) derived from transgenic mice that moderately over-express a TAP-epitope tagged human 14-3-3ζ molecule (TAP-14-3-3ζ) (4,31) (Figure 1A). This approach was chosen to circumvent the variability in the expression of transiently expressed proteins and increased specificity of protein purification with epitope-tagged proteins (32,33). Differentiation of MEFs was induced with an established adipogenic cocktail (insulin, dexamethasone, and IBMX), supplemented with rosiglitazone (Figure 1A,B). Differentiation into adipocytes was confirmed by Oil Red-O staining and *Pparg* mRNA expression (Figure 1B,C).

**Figure 1:**
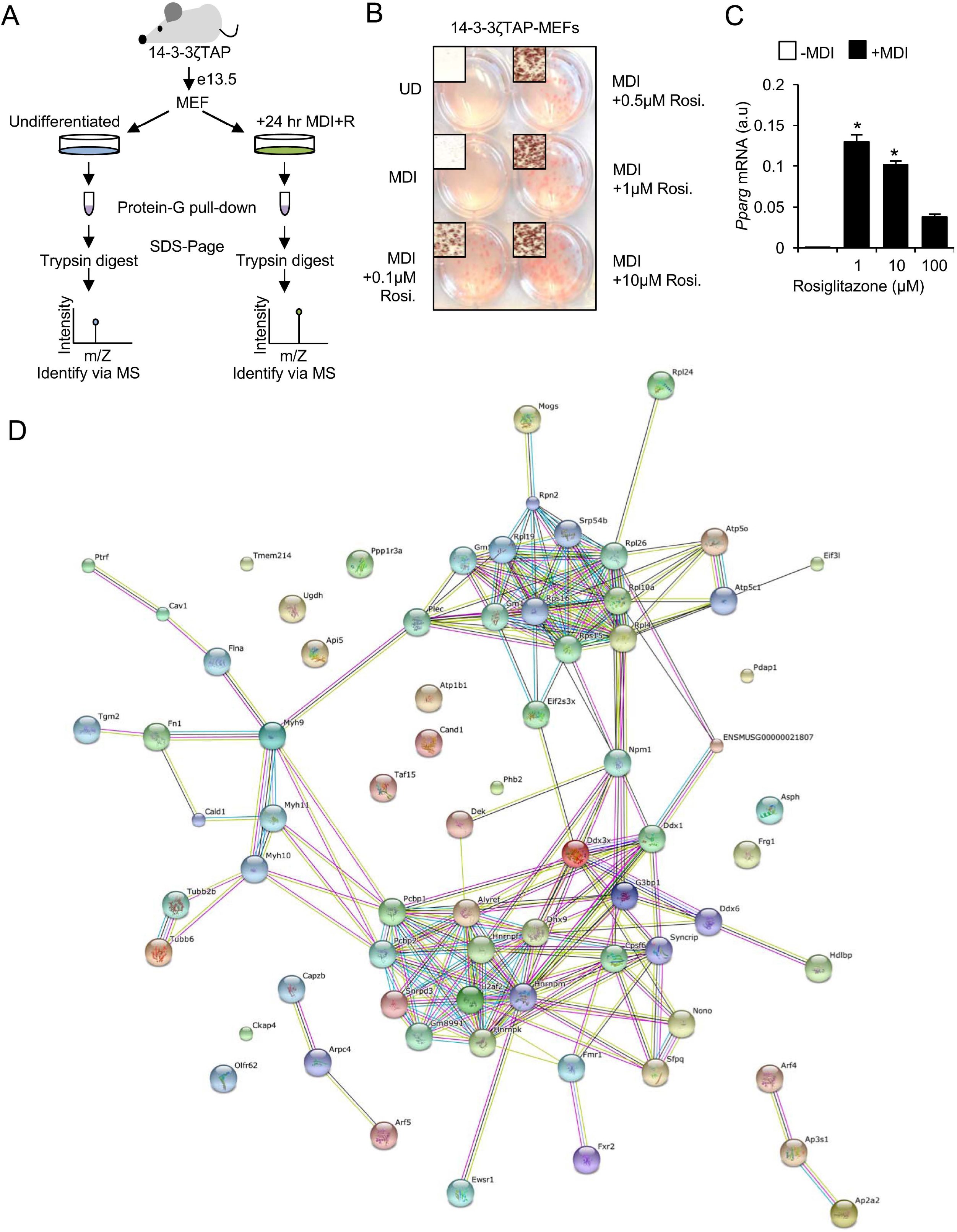
Generation of TAP-14-3-3ζ mouse embryonic fibroblasts (MEFs) to elucidate the 14-3-3ζ interactome. **(A)** Schematic over-view of generation and use of TAP-14-3-3o MEFs to determine the 14-3-3ζ interactome during adipogenesis. **(B,C)** Verification of TAP-14-3-3ζ MEF adipogenesis by Oil Red-O incorporation, 7 days after induction (B), or *Pparg* mRNA expression by quantitative PCR (C), 2 days following induction (representative of n=4 independent experiments, *: p<0.05). **(D)** String-db (24) was used to visualize and cluster proteins according to their biological function, resulting in three distinct clusters: RNA splicing/processing factors, components of the ribosomal complex, and components of actin/tubulin network.

### 3.2 Differentiation of TAP-14-3-3ζ MEFs results in distinct changes in the interactome of 14-3-3ζ

Although we previously identified the hedgehog signaling effector, Gli3, as a downstream regulator of 14-3-3ζ-dependent adipogenesis (4), we hypothesized that 14-3-3ζ may control other parallel processes underlying adipocyte differentiation. This is due in part to the large, diverse interactomes of 14-3-3 proteins (8,16–18). Thus, we utilized affinity proteomics to identify interacting proteins that associate with 14-3-3ζ during adipocyte differentiation (Figure 1A). The interactome of 14-3-3ζ at 24 hours post-induction was examined because key signaling events underlying murine adipocyte differentiation occur during the first 24-48 hours (2,4,34). Over 100 proteins were identified by mass spectrometry as 14-3-3ζ interacting proteins (Table 1). Of these proteins, 56 have not been previously reported to interact with any member of the 14-3-3 protein family (Table 2) (17). 14-3-3ζ itself was found equally enriched in both samples, demonstrating equal pull-down efficiency (data not shown). An enrichment of differentiation-dependent 14-3-3ζ-interacting proteins associated with RNA splicing, translation, protein transport, and nucleic acid transport were detected using gene ontology to define their biological processes (23,24) (Table 3). Thus, these proteomic data demonstrate the dynamic nature of the 14-3-3ζ interactome and suggest that 14-3-3ζ may regulate multiple processes required for adipocyte differentiation through its interactions.

**Table 1:**
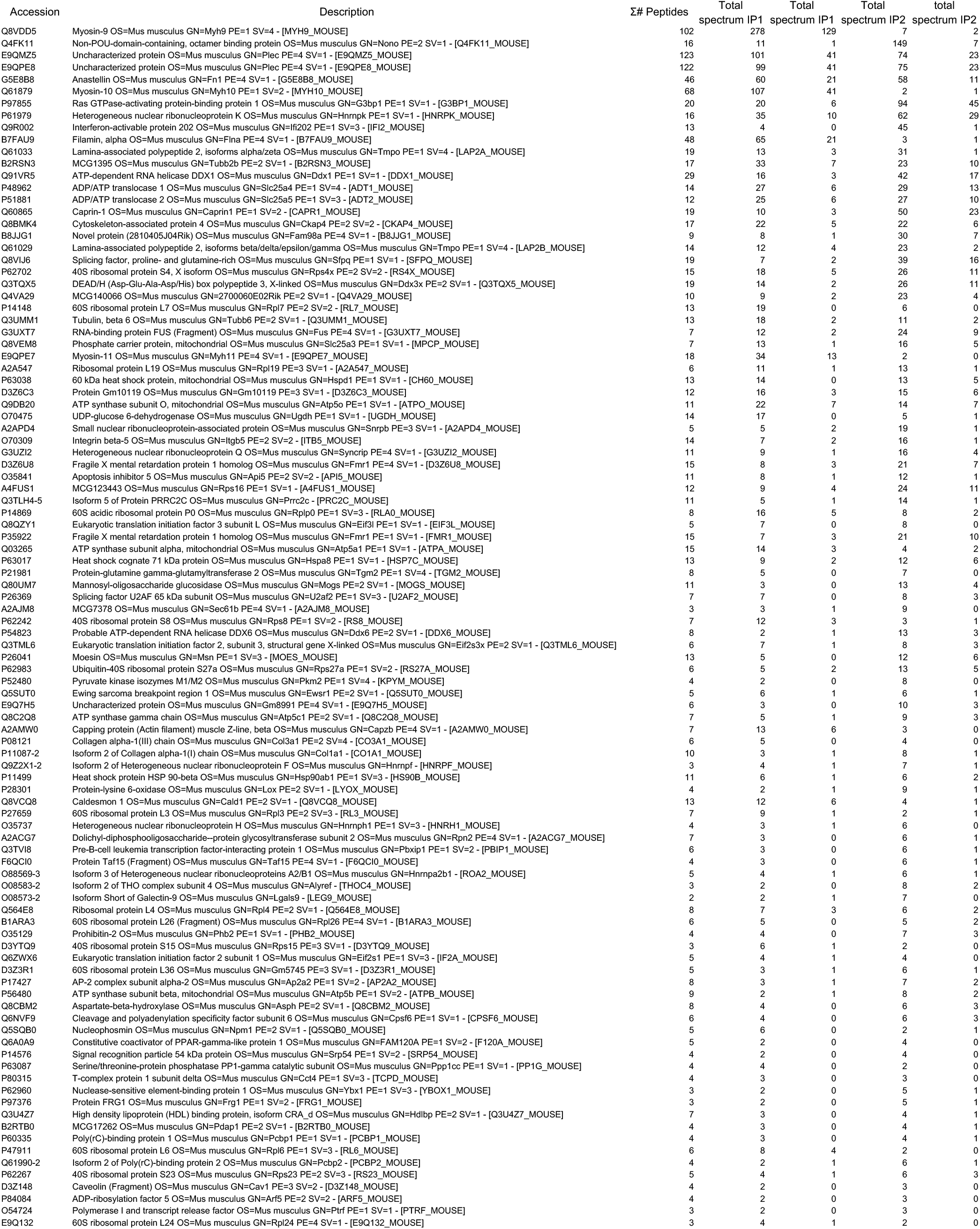
Proteins with at least 2 unique peptides with a total spectral count in differentiated cell >=2 in comparison to undifferentiated cells.

**Table 2:**
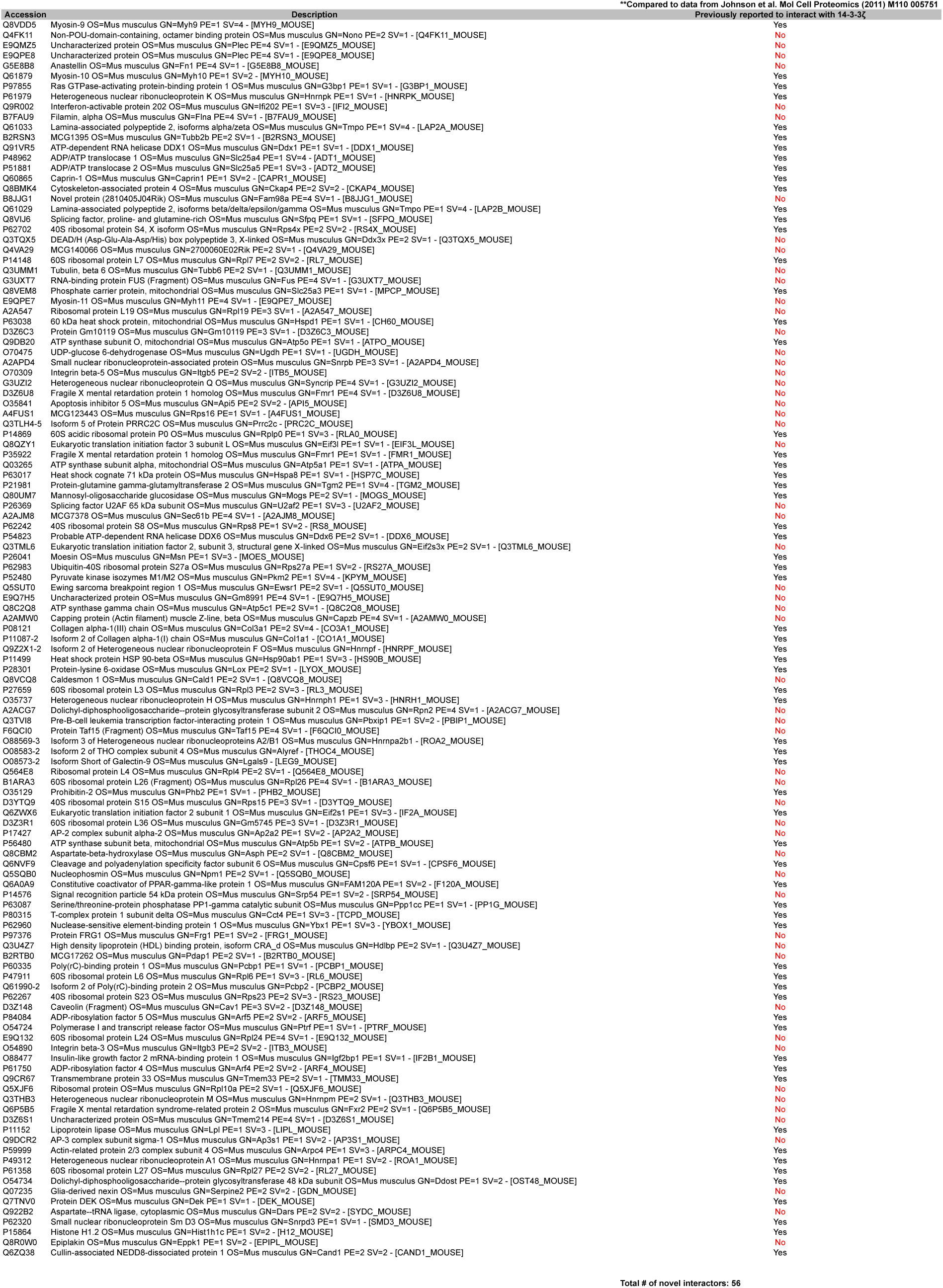
Identification of novel interactors with 14-3-3 proteins.

** Compared to data from Johnson et al. Mol Cell Proteomics (2011) M110 005751

**Table 3:** Gene ontology classification of proteomic hits by biological process.

**Annotation Cluster 1**
| Term |  | p-value | Benjamini | FDR |
| --- | --- | --- | --- | --- |
| GO:0006397 | mRNA processing | 2.80E-12 | 1.18E-09 | 4.33E-09 |
| GO:0008380 | RNA splicing | 8.22E-12 | 2.31E-09 | 1.27E-08 |
| GO:0016071 | mRNA metabolic process | 2.70E-11 | 5.71E-09 | 4.18E-08 |
| GO:0006396 | RNA processing | 1.09E-09 | 1.84E-07 | 1.68E-06 |

**Annotation Cluster 2**
| Term |  | p-value | Benjamini | FDR |
| --- | --- | --- | --- | --- |
| GO:0065003 | macromolecular complex assembly | 1.23E-04 | 0.01719154 | 0.19018503 |
| GO:0006461 | protein complex assembly | 1.87E-04 | 0.01956598 | 0.28880909 |
| GO:0070271 | protein complex biogenesis | 1.87E-04 | 0.01956598 | 0.28880909 |
| GO:0043933 | macromolecular complex subunit organization | 2.40E-04 | 0.02229116 | 0.37052613 |
| GO:0034621 | cellular macromolecular complex subunit organization | 0.001634086 | 0.10877737 | 2.49672826 |
| GO:0034622 | cellular macromolecular complex assembly | 0.004135111 | 0.20818502 | 6.20540124 |
| GO:0043623 | cellular protein complex assembly | 0.00690653 | 0.26524962 | 10.1607298 |
| GO:0051258 | protein polymerization | 0.031762743 | 0.57358582 | 39.2881889 |

**Annotation Cluster 3**
| Term |  | p-value | Benjamini | FDR |
| --- | --- | --- | --- | --- |
| GO:0015931 | nucleobase, nucleoside, nucleotide and nucleic acid transport | 1.65E-04 | 0.01976438 | 0.25533903 |
| GO:0051236 | establishment of RNA localization | 0.001160158 | 0.09343294 | 1.77868077 |
| GO:0050658 | RNA transport | 0.001160158 | 0.09343294 | 1.77868077 |
| GO:0050657 | nucleic acid transport | 0.001160158 | 0.09343294 | 1.77868077 |
| GO:0006403 | RNA localization | 0.001227278 | 0.09002225 | 1.88067318 |

**Annotation Cluster 4**
| Term |  | p-value | Benjamini | FDR |
| --- | --- | --- | --- | --- |
| GO:0015986 | ATP synthesis coupled proton transport | 0.002165467 | 0.13143078 | 3.29598278 |
| GO:0015985 | energy coupled proton transport, down electrochemical gradient | 0.002165467 | 0.13143078 | 3.29598278 |
| GO:0034220 | ion transmembrane transport | 0.00311983 | 0.17188096 | 4.71608221 |
| GO:0015992 | proton transport | 0.005711473 | 0.26103289 | 8.47469156 |
| GO:0006818 | hydrogen transport | 0.006023853 | 0.24696352 | 8.91824507 |
| GO:0006119 | oxidative phosphorylation | 0.007021577 | 0.25748192 | 10.3215008 |
| GO:0006754 | ATP biosynthetic process | 0.019701699 | 0.53433002 | 26.4817286 |
| GO:0046034 | ATP metabolic process | 0.025117659 | 0.59165663 | 32.5166079 |
| GO:0009201 | ribonucleoside triphosphate biosynthetic process | 0.027334987 | 0.59373718 | 34.8509636 |
| GO:0009206 | purine ribonucleoside triphosphate biosynthetic process | 0.027334987 | 0.59373718 | 34.8509636 |
| GO:0009145 | purine nucleoside triphosphate biosynthetic process | 0.028096637 | 0.59012787 | 35.6352295 |
| GO:0009142 | nucleoside triphosphate biosynthetic process | 0.02886954 | 0.5741125 | 36.4220479 |
| GO:0009205 | purine ribonucleoside triphosphate metabolic process | 0.033742375 | 0.58476996 | 41.1791708 |
| GO:0009199 | ribonucleoside triphosphate metabolic process | 0.034593574 | 0.58312761 | 41.9751939 |
| GO:0006091 | generation of precursor metabolites and energy | 0.03610049 | 0.58839461 | 43.3597731 |
| GO:0009144 | purine nucleoside triphosphate metabolic process | 0.038109124 | 0.58825354 | 45.1573304 |
| GO:0009152 | purine ribonucleotide biosynthetic process | 0.039015563 | 0.58726975 | 45.9509181 |
| GO:0055085 | transmembrane transport | 0.042081945 | 0.60604395 | 48.5566327 |
| GO:0009260 | ribonucleotide biosynthetic process | 0.042750615 | 0.59362044 | 49.1090183 |
| GO:0009141 | nucleoside triphosphate metabolic process | 0.046658895 | 0.60897427 | 52.228246 |

**Annotation Cluster 5**
| Term |  | p-value | Benjamini | FDR |
| --- | --- | --- | --- | --- |
| GO:0001568 | blood vessel development | 0.028163983 | 0.57774561 | 35.7041482 |
| GO:0001944 | vasculature development | 0.030823763 | 0.58598984 | 38.3715132 |

Enrichment Score: 1.1516193992206216

### 3.3 Regulation of mRNA processing by 14-3-3ζ

Using our previous transcriptomic analysis of differentiating 3T3-L1 cells (26), we re-analyzed the effects of differentiation and 14-3-3ζ depletion on RNA processing. Differential exon usage (DEXSeq) was used as a surrogate measure of alternative splicing of mRNA (Figure 2A) (25). Any changes in splice variant levels were not due to global effects of 14-3-3ζ depletion on RNA transcription, as no gross differences in the incorporation of a uracil analog were detected (Figure 2B). Comparison of genes that displayed differential exon usage at 24 and 48 hours post differentiation revealed that 163 and 172 genes, respectively, that were unique to each time point (Figure 2C). Gene ontology analysis revealed that at each time point, distinct groups of genes were alternatively spliced (Table 4). The use of this approach to detect genes with differential exon usage was validated by the ability to detect *Pparg* variants after 48 hours of differentiation (Figure S1) (35). The effect of 14-3-3ζ depletion was assessed at each time point, and 78, 37, and 36 genes were affected following 14-3-3ζ knockdown at 0, 24, and 48 hours, respectively, after the induction of differentiation (Figure 2D). However, only in undifferentiated 3T3-L1 cells could enrichments in genes associated with macromolecular complex assembly (GO:0065003, p=3.44 × 10^−3^), macromolecular complex subunit organization (GO:0043933, p= 7.56 × 10^−4^), and regulation of biological quality (GO:0065008, p=9.51 × 10^−3^) be detected by gene ontology analysis. Collectively, these data demonstrate that adipogenesis promotes the alternative splicing of genes and this process can be influenced by 14-3-3ζ.

**Figure 2:**
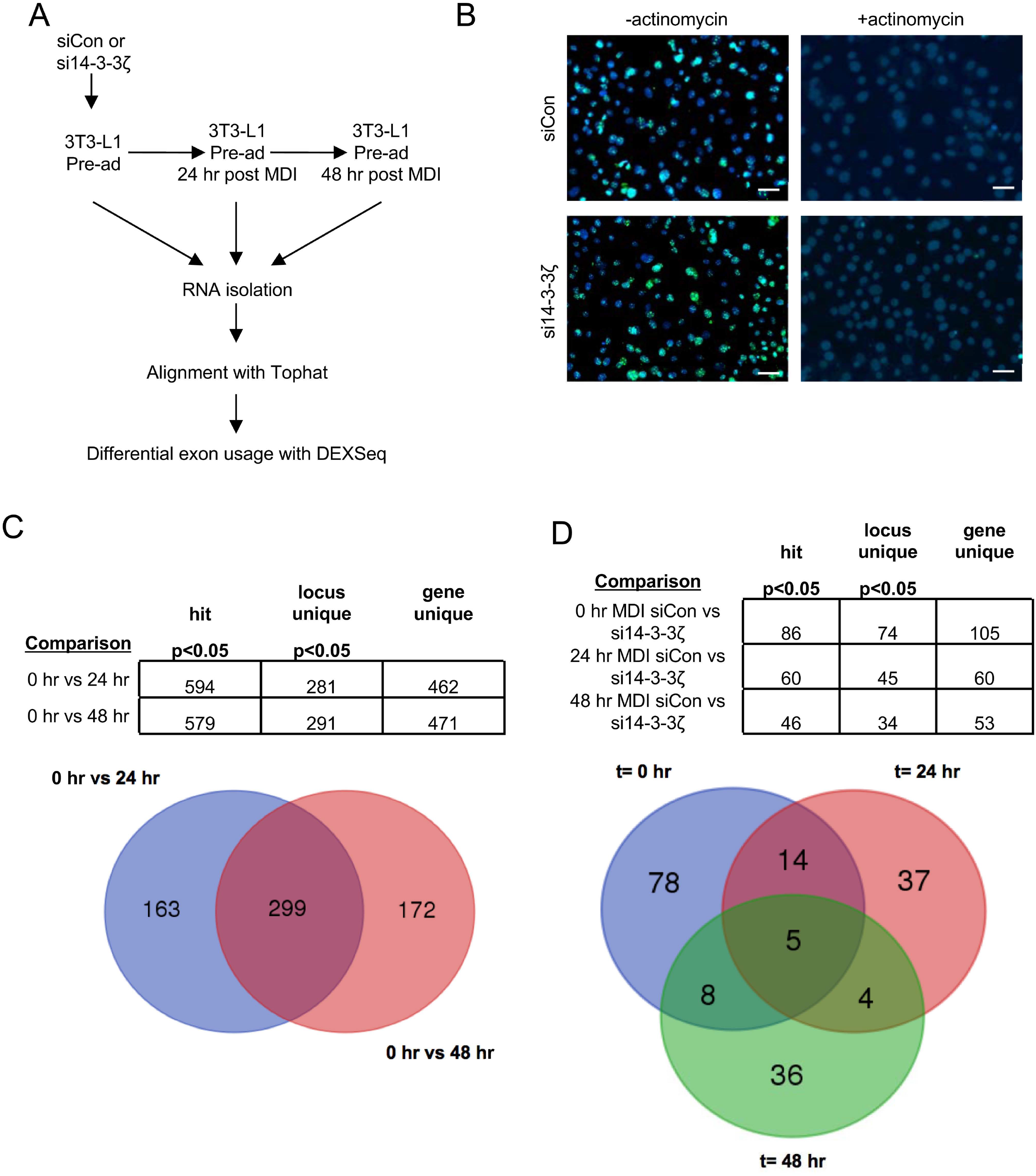
Induction of differentiation or depletion of 14-3-3ζ in 3T3-L1 cells promotes alternative splicing of mRNA. **(A)** Differential exon usage of genes involved in adipogenesis was compared in control or 14-3-3ζ-depleted 3T3-L1 cells undergoing adipocyte differentiation. Transcriptomic data was aligned via TopHat and subsequently subjected to DEXSeq analysis to measure differential exon usage. **(B)** To rule out an effect of 14-3-3ζ depletion on global RNA transcription, control (siCon) or 14-3-3ζ depleted cells (si14-3-3ζ) were incubated with 5-ethynyl uridine (EU), followed by Click-iT chemistry to detect newly synthesized RNA (scale bar= 10 μm; representative of n=4 experiments). **(C,D)** Comparison of genes exhibiting differential exon usage in control cells 0, 24, and 48 hours after differentiation (C) or control or 14-3-3ζ depleted cells at each time point (D). The overlapping regions of each Venn diagram denote genes that are common to each condition or treatment.

**Table 4.**
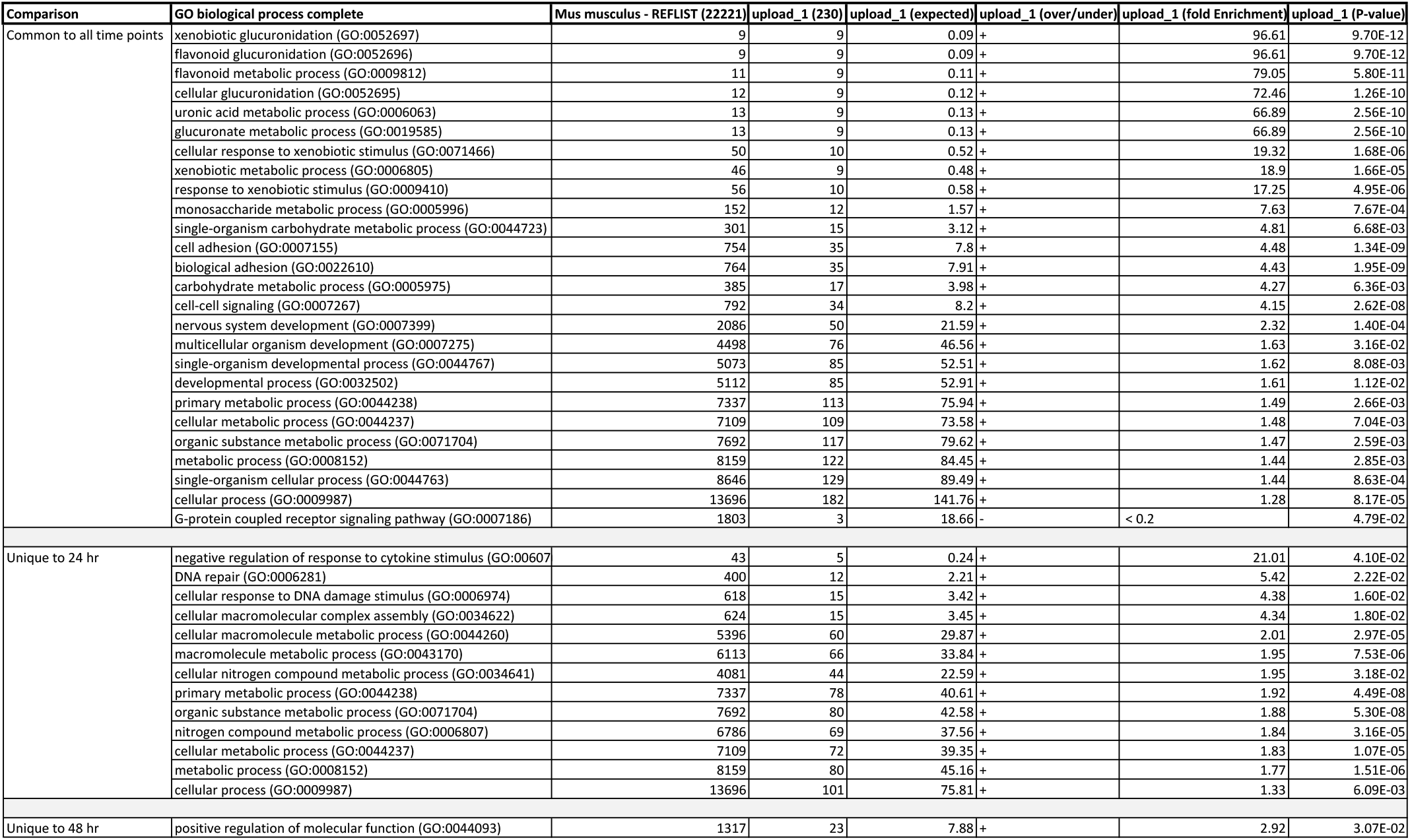
Analysis of common and unique genes during the first 48 hrs of 3T3-L1 adipogenesis.

### 3.4 Identification of known and novel regulators of adipocyte differentiation

Within the 14-3-3ζ interactome, we were able to detect proteins with known roles in adipogenesis, such as Ptrf/Cavin1 and prohibitin-2 (Phb2) (36–40) and confirmed their roles in adipocyte differentiation (Figure 3). This confirmed that known regulators of adipogenesis can be detected within the 14-3-3ζ interactome and suggested the possibility that novel factors could be identified. Additional proteins in the 14-3-3ζ interactome, such as Fragile-X mental retardation protein-1 (Fmr1) and Rpn2, were also examined for their roles in adipogenesis, as they have previously been shown to be associated with obesity or weight gain (41,42). However, siRNA-mediated knockdown of either protein had no effect on 3T3-L1 differentiation, indicating that these proteins are not required for adipogenesis (Figure 3A-D), at least in this *in vitro* model system.

**Figure 3:**
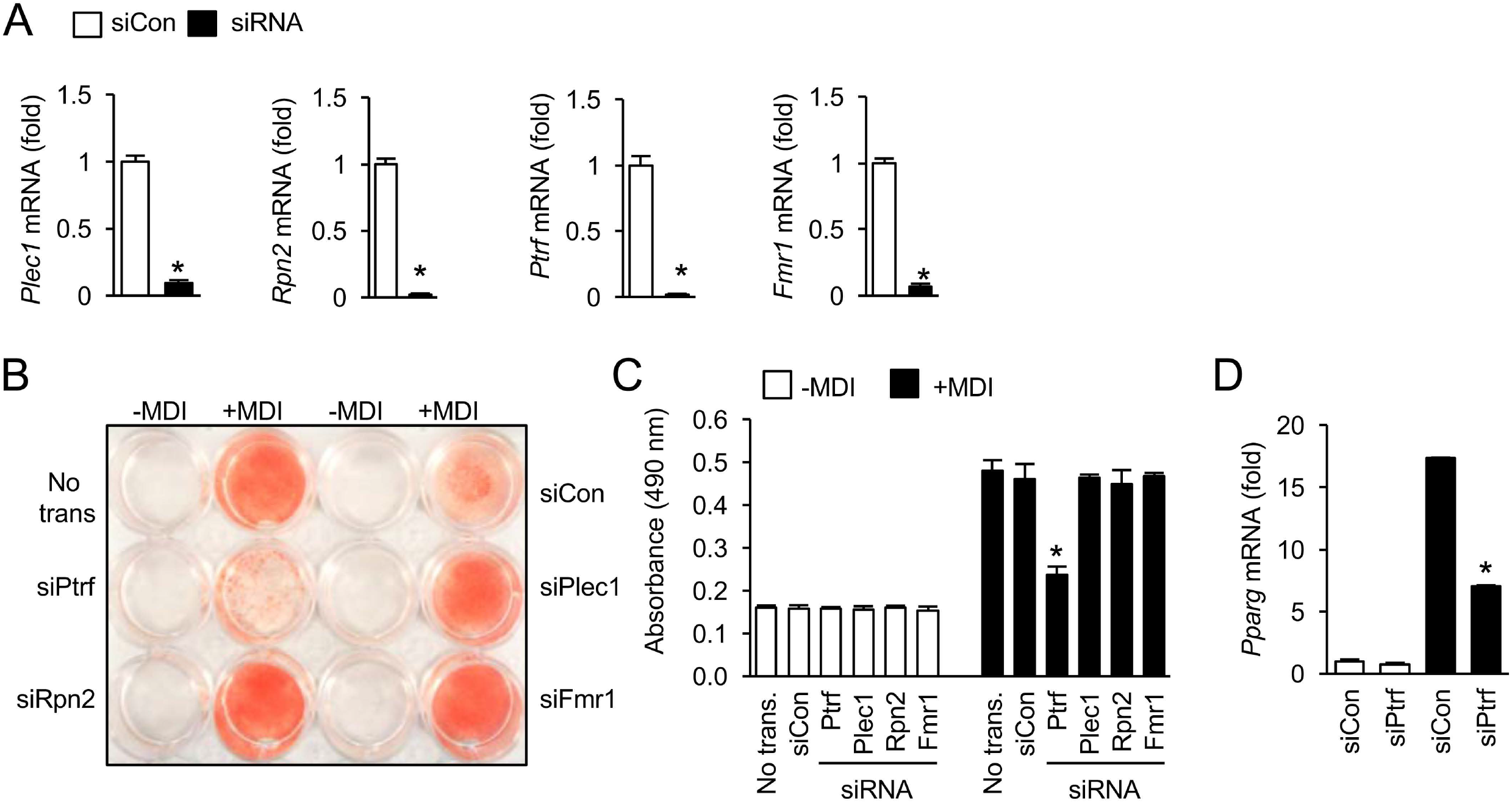
Known regulators of adipogenesis can be found within the 14-3-3ζ interactome. **(A)** 3T3-L1 cells were transfected with a control siRNA (siCon) or siRNA against target mRNA, and knockdown efficiency was measured by quantitative PCR (n=4 per group, *: p<0.05). **(B, C)** Transient knockdown by siRNA of previously identified regulators of adipogenesis or those associated with the development of obesity was used to examine their contributions to adipocyte differentiation, as assessed by visualization of Oil Red-O incorporation (B), absorbance (490 nm, C) or *Pparg* mRNA expression (D) (n=4 per group, *:p<0.05 when compared to siCon-transfected differentiated cells).

As proteins associated with RNA processing and splicing were highly enriched during differentiation (Table 3), we sought to examine contribution of RNA splicing factors to adipogenesis. Using siRNA in 3T3-L1 pre-adipocytes, 8 splicing factors, which were identified in our proteomic analysis of the 14-3-3ζ interactome (Table 1), were screened for their roles in 3T3-L1 adipogenesis. They were chosen by the number of connections exhibited within each cluster of proteins (Figure 1D) (24). Of note, mRNA levels of the chosen splicing factors were generally unaffected by knockdown of 14-3-3ζ; however, some splicing factors were influenced by differentiation (Figure S2) (26). Transient knockdown of Ddx6, Sfpq, Hnrnpf, or Hnrnpk was sufficient to impair 3T3-L1 differentiation, as assessed by Oil Red-O incorporation (Figure 4). Closely related proteins with similar roles, such as Ddx1, Nono, Hnrnpm, and Syncrip/Hnrnpq were not required for 3T3-L1 adipogenesis (Figure 4B, C). Knockdown of Ddx6 or Hnrnpk by siRNA did not have an effect of *Pparg*, which suggests that these factors act downstream of the mRNA expression of this master transcription factor (Figure 4D). Other pro- or anti-adipogenic genes are alternatively spliced during adipocyte differentiation. For example, *Lpin1* mRNA is spliced to generate Lipin-1α and Lipin-1 β, which have differential roles on adipogenesis (28). To examine the effect of depletion of 14-3-3ζ, Hnrnpf, Ddx6, Hnrnpk, and Sfpq on *Lpin1* splicing, 3T3-L1 cells were transiently transfected with siRNA, followed by the induction of differentiation. Gene silencing of all target genes was found to prevent the generation of the *Lpin-1α* variant during differentiation (Figure 4E, F). Collectively, these findings demonstrate that novel regulators of adipogenesis can be identified within the interactome of 14-3-3ζ and highlight the involvement of 14-3-3ζ in regulating the alternative splicing of mRNA.

**Figure 4:**
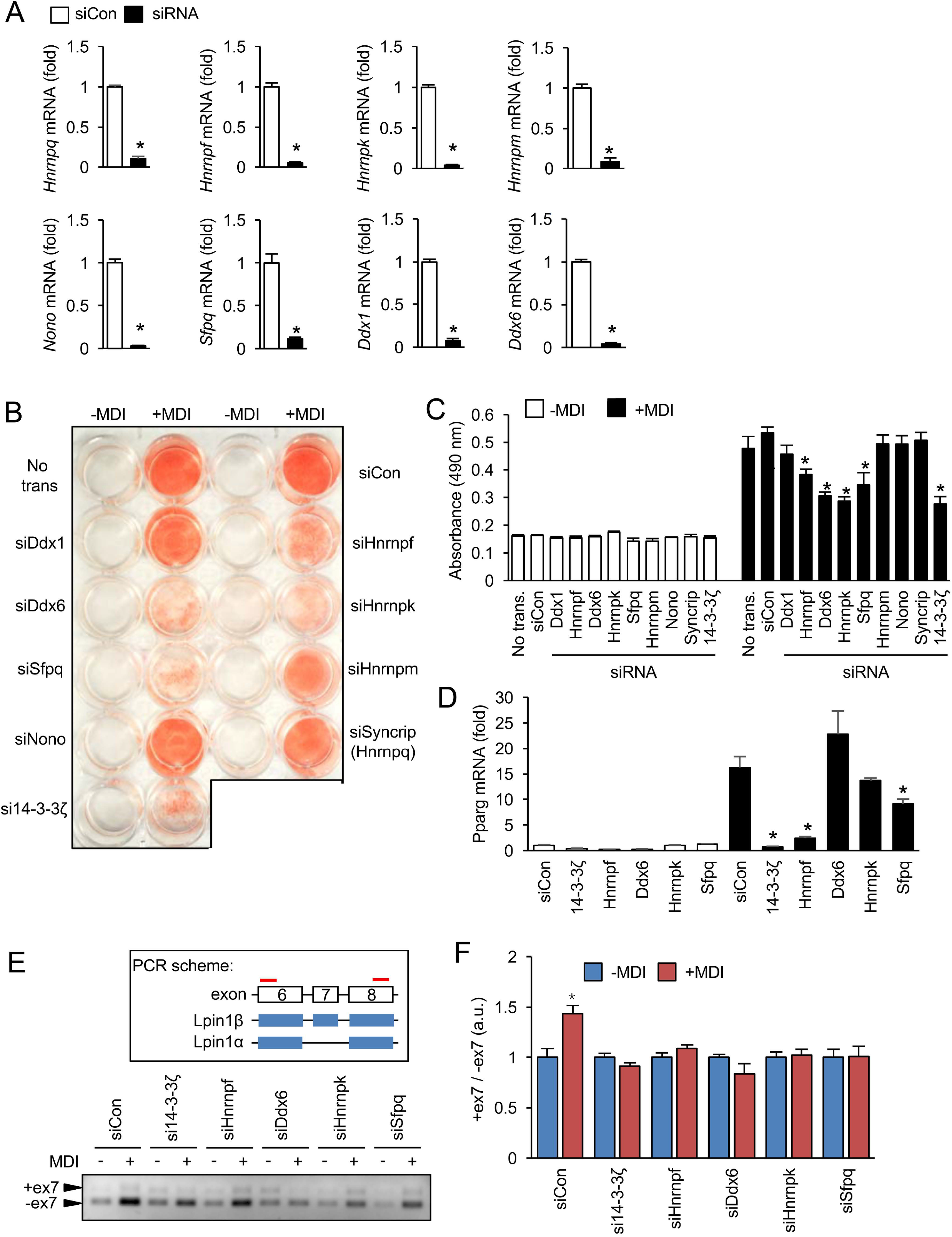
RNA splicing proteins are required for 3T3-L1 adipogenesis. **(A)** 3T3-L1 cells were transfected with a control siRNA (siCon) or siRNA against target mRNA, and knockdown efficiency was measured by quantitative PCR (n=4 per group, *: p<0.05). **(B-D)** Transient knockdown by siRNA was used to examine the contributions of RNA splicing factors to adipocyte differentiation, as assessed by visualization of Oil Red-O incorporation (B), absorbance (490 nm, C) or *Pparg* mRNA expression (D) (n=4 per group, *:p<0.05 when compared to siCon-transfected differentiated cells). **(E)** 3T3-L1 cells were transfected with siRNAs against various targets, followed by induction of differentiation (+MDI) for 48 hours, and total RNA was isolated. Following RT-PCR for *Lpin1*, products were resolved on a 1% acrylamide gel (Inset: Schematic diagram of the RT-PCR-based approach to detect alternative spliced isoforms of *Lpin1*. Red bars denote primers used to detected the inclusion or exclusion of exon 7) (representative of n=4 independent experiments) **(F)** Densitometric analysis of *Lpin1* PCR products from panel E (*: p<0.05 when compared to undifferentiated (-MDI) 3T3-L1 cells).

## 4. Discussion

In the present study, affinity proteomics was used to determine how adipogenesis influences the interactome of 14-3-3ζ. Surprisingly, the interactome was dynamic, as differentiation altered the landscape of proteins that interact with 14-3-3ζ. This approach also permitted the identification of known adipogenic factors within the 14-3-3ζ interactome and revealed novel proteins that are required for adipocyte differentiation. An enrichment of proteins associated with RNA processing and splicing were detected, and the novel contributions of RNA splicing factors, such as Hnrnpf, Ddx6, and Sfpq, in adipogenesis were identified. The usefulness of this approach was also evident in the ability to identify process that may be regulated by 14-3-3ζ during adipocyte differentiation.

We previously identified an essential function of the hedgehog signaling effector Gli3 in 14-3-3ζ-regulated adipocyte differentiation (4). However, due to the large, diverse interactome of 14-3-3 proteins (10,13,16,17), we hypothesized that it is unlikely that one protein would be solely responsible for 14-3-3ζ-mediated adipogenesis. It is known that the interactomes of 14-3-3 proteins are dynamic and change in response to various stimuli (11–13,19). The functional significance of such changes in the interactome is not clear, but it suggests that 14-3-3 proteins may be regulating biological processes critical for adipocyte development through their interactions. Using a gene onotology-based approach, we found that the 14-3-3ζ interactome is enriched with proteins involved in RNA binding and splicing during differentiation and confirms its contribution to the alternative splicing of mRNAs. As over 100 proteins were found to be unique to the14-3-3ζ interactome during adipocyte differentiation, it suggests that 14-3-3ζ could also regulate other cellular processes required for adipocyte development. For example, we detected an interaction of 14-3-3ζ with the mitochondrial regulator, Prohibitin-2 (Phb2), which others have shown to be essential for the expansion of mitochondria mass and mitochondrial function during adipogenesis (36–38). Further in-depth studies are required to assess whether 14-3-3ζ has regulatory roles in mitochondrial dynamics, but when taken together, it demonstrates the possibility of examining the contributions of interacting partners to reveal novel biological processes required for adipocyte differentiation.

Through the use of a functional siRNA screen, we identified novel roles of various RNA splicing factors, namely Hnrpnf, Hnrnpk, Ddx6, and Sfpq, in adipocyte differentiation. Sfpq belongs to the Drosophila behavior/human splicing (DHBS) protein family and is required for transcriptional regulation (43,44). Although a recent study by Wang and colleagues found no effect of forced overexpression of Nono and Sfpq on adipogenesis (45), we report that Sfpq depletion impairs adipocyte differentiation. DHBS proteins may exhibit redundant, compensatory functions (46), but given that only Sfpq depletion impaired 3T3-L1 adipogenesis, it suggests specific protein-protein or protein-nucleic acid interactions occur may with each DHBS member in the context of differentiation (44). We were also able to detect novel adipogenic roles of Hnrnpf and Hnrnpk, members of the heterogeneous nuclear ribonucleoproteins (Hnrnps) which facilitate mRNA splicing (47,48). Alternative splicing of mRNA is critical for maintaining genetic diversity and cell identity, in addition to the expression of key factors required for differentiation (49,50). Specific to adipogenesis, differential promoter usage and alternative splicing are required for the expression of the canonical adipogenic transcription factor PparY (51–53). Other regulatory factors are also formed from alternative splicing, including nCOR1 and Lipin1 (54,55). Future studies are required to determine whether 14-3-3ζ directly binds to these splicing factors and how it regulates their splicing activity to generate essential adipogenic factors.

Protein abundance of 14-3-3ζ and other isoforms is increased in visceral adipose tissue from obese individuals (56,57), and we have previously reported that systemic over-expression of 14-3-3ζ in mice is sufficient to potentiate weight gain and fat mass in mice fed a high-fat diet (4). With respect to the pancreatic β-cell, single cell transcriptomic analysis revealed higher mRNA expression of *YWHAZ* in β-cells from subjects with type 2 diabetes (58), and we have found that systemic over-expression of 14-3-3ζ was sufficient to reduce β-cell secretory function in mice (9). The exact mechanisms owing to how changes in 14-3-3ζ function affects the development of obesity or β-cell dysfunction are not known, but In-depth examination of the interactome in the context of both conditions may yield novel biological insight as to how 14-3-3ζ influences their development. This approach has already been useful in understanding how changes in 14-3-3ε or 14-3-3σ expression can lead to the development of various forms of cancer and the identification of novel therapeutic targets (19,59–61).

In conclusion, this study provides compelling evidence demonstrating the usefulness of elucidating the interactome of 14-3-3ζ as a means to identify novel factors required for adipogenesis. Additionally, a systematic investigation of interacting partners may also provide insight as to which physiological processes are essential for 14-3-3ζ-mediated adipocyte differentiation. Lastly, deciphering how various disease states influence the interactome of 14-3-3 proteins may also aid in the discovery of novel therapeutic targets for the treatment of chronic diseases, such as obesity and type 2 diabetes.

## 5. Acknowledgements

This work was supported by a CIHR Project grant (PJT-153144) to GEL. Some of this work was initiated by GEL when he was a JDRF- and Canadian Diabetes Association-supported postdoctoral fellow in the lab of Dr. James D. Johnson (University of British Columbia, Vancouver, BC, Canada). The authors would like to thank François Harvey in the Bioinformatics platform at the CRCHUM for bioinformatics support, as well as Dr. Johnson for critical reading of this manuscript.

## 6. Authors contribution

Y.M performed experiments, analyzed data, and wrote and reviewed the manuscript. MS and NNF performed experiments and analyzed data. TM designed parts of the study and reviewed the manuscript. GEL performed experiments, analyzed data, wrote the manuscript, and is responsible for the integrity of this work.

---

**Supplemental figure 1:**
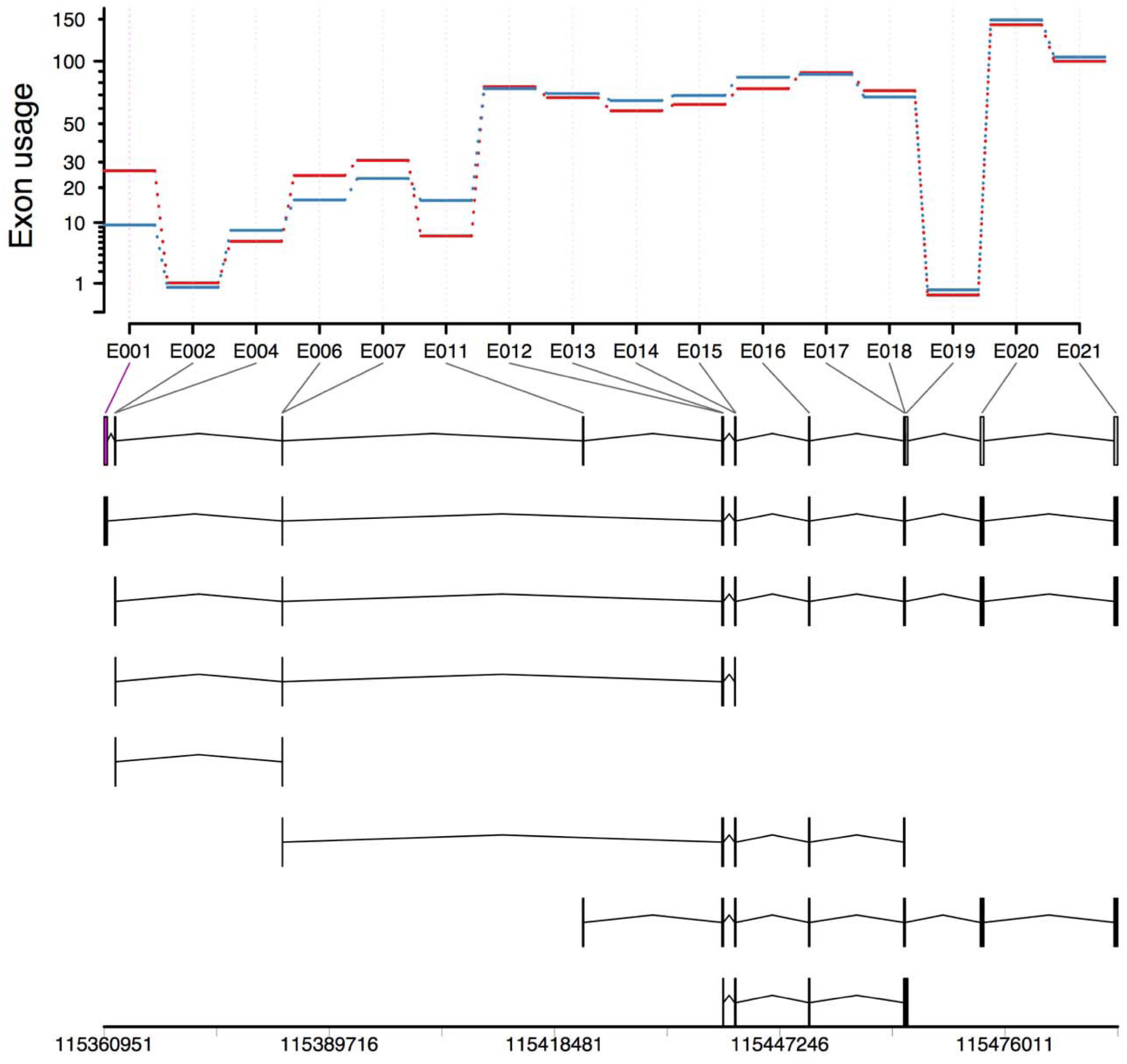
*Pparg* exhibits differential exon usage during 3T3-L1 adipocyte differentiation. To confirm the ability of DEXSeq to detect genes exhibiting significant differential exon usage, transcriptomic data from undifferentiated and differentiating 3T3-L1 cells (48 hours post induction) were analyzed, and isoforms with differential use of exon 1 could be detected (n=4 per group).

**Supplemental figure 2:**
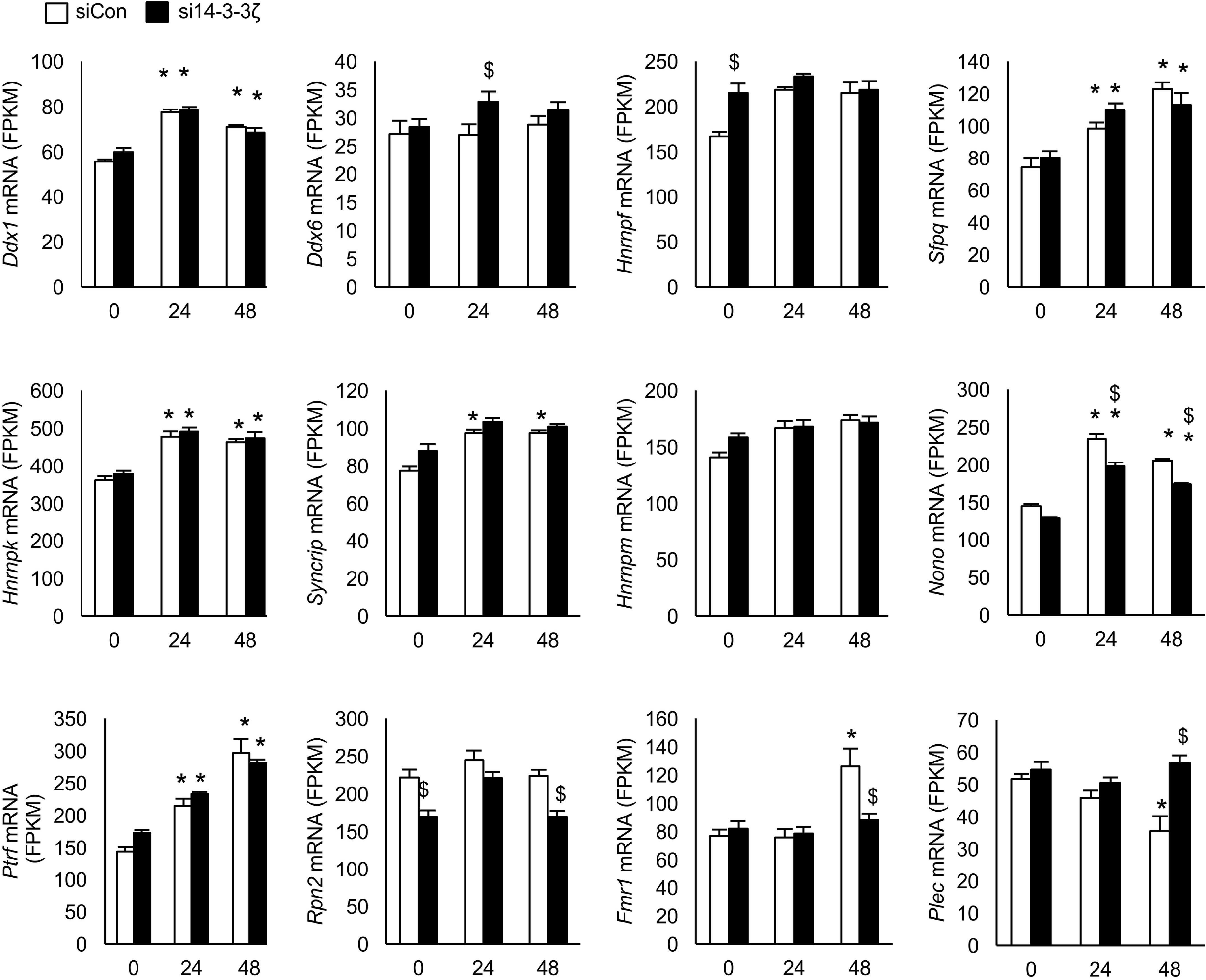
Expression of candidate proteins from the 14-3-3ζ proteomic screen are largely unaffected by depletion of 14-3-3ζ. Transcriptomic data [GSE60745] (26) from 3T3-L1 cells transfected with control (siCon) or siRNA against 14-3-3ζ (si 14-3-3ζ), followed by differentiation with an adipogenic cocktail (MDI) for up to 48 hours. The dataset was queried for expression profiles of 14-3-3ζ interacting partners that will be tested for their adipogenic contributions (Figure 3). (n=4 per group, *: p<0.05 when compared to t=0, $: p<0.05 when compared to siCon at respective time point).

